# Advancing Protein-DNA Binding Site Prediction: Integrating Sequence Models and Machine Learning Classifiers

**DOI:** 10.1101/2023.08.23.554389

**Authors:** Taslim Murad, Prakash Chourasia, Sarwan Ali, Murray Patterson

## Abstract

Predicting protein-DNA binding sites is a challenging computational problem in the field of bioinformatics. Identifying the specific residues where proteins bind to DNA is of paramount importance, as it enables the modeling of their interactions and facilitates downstream studies. Nevertheless, the development of accurate and efficient computational methods for this task remains a persistent challenge. Accurate prediction of protein-DNA binding sites has far-reaching implications for understanding molecular mechanisms, disease processes, drug discovery, and synthetic biology applications. It helps bridge the gap between genomics and functional biology, enabling researchers to uncover the intricacies of cellular processes and advance our knowledge of the biological world. The method used to predict DNA binding residues in this study is a potent combination of conventional bioinformatics tools, protein language models, and cutting-edge machine learning and deep learning classifiers. On a dataset of protein-DNA binding sites, our model is meticulously trained, and it is then rigorously examined using several experiments. As indicated by higher predictive behavior with AUC values on two benchmark datasets, the results show superior performance when compared to existing models. The suggested model has a strong capacity for generalization and shows specificity for DNA-binding sites. We further demonstrated the adaptability of our model as a universal framework for binding site prediction by training it on a variety of protein-ligand binding site datasets. In conclusion, our innovative approach for predicting protein-DNA binding residues holds great promise in advancing our understanding of molecular interactions, thus paving the way for several groundbreaking applications in the field of molecular biology and genetics. Our approach demonstrated efficacy and versatility underscore its potential for driving transformative discoveries in biomolecular research.

## 1 Introduction

Protein-DNA binding site prediction is an essential area of research with significant implications in various fields, including molecular biology, genetics, drug discovery, and synthetic biology. Accurate prediction of protein-DNA binding sites has far-reaching potential, leading to groundbreaking biotechnology and drug design applications [50] along with several other applications including understanding gene regulation [30], functional annotation of genomes [28], cancer research for designing targeted therapies [40], evolutionary studies [18], genetic engineering, and accelerating drug discovery [47]. Protein-DNA interactions are fundamental to many biological processes, such as DNA replication [8], transcription [7], repair, and recombination [29]. Accurate prediction of binding sites aids in deciphering the molecular mechanisms behind these processes, shedding light on the intricate workings of the cell. It plays a crucial role in regulating gene expression by binding to specific sites on DNA. Predicting these binding sites helps researchers understand how genes are turned on or off, which is essential for understanding normal development, disease processes, and various cellular responses. It also can help identify potential drug targets for diseases like cancer and genetic disorders. Some research applies protein-DNA binding site prediction to specific diseases, such as cancer [48]. By identifying altered binding sites in disease-related genes, researchers aim to uncover novel therapeutic targets. Designing drugs that interfere with these interactions could offer new therapeutic strategies. With the advent of high-throughput sequencing technologies, vast amounts of genomic data are generated. The binding site prediction for protein and DNA aids in interpreting this data by identifying regions that are likely to be functionally important. Protein-DNA binding sites in genomic analyses provide a foundational understanding of the functional elements within a genome and their roles in various biological processes. It bridges the gap between genomic sequences and biological functions, enabling researchers to unravel the complexities of gene regulation and molecular interactions.

Many recent studies leverage machine learning and deep learning techniques to predict protein-DNA binding sites [46]. These methods often involve training models on large datasets of known binding sites and using them to predict binding locations in genomic sequences. Researchers focus on identifying relevant features or descriptors that can capture the characteristics of protein-DNA interactions [15]. These features might include sequence motifs, physicochemical properties, and structural information of DNA and protein molecules [14]. Some studies integrate various types of omics data, such as genomics, transcriptomics, and proteomics, to improve the accuracy of binding site predictions. This integration allows for a more comprehensive understanding of the regulatory landscape. Evolutionarily conserved regions are often indicative of functional importance [35]. Research explores the use of conservation scores and comparative genomics approaches to enhance the accuracy of binding site predictions.

Incorporating 3D structural information of protein-DNA complexes helps refine binding site predictions by considering the physical interactions between the molecules. This includes techniques such as molecular docking and structural modeling. Deep learning models, such as convolutional neural networks (CNNs) and recurrent neural networks (RNNs), are applied to sequence data to capture complex patterns in DNA and protein sequences, improving binding site prediction accuracy. Transfer learning techniques involve pretraining models on related tasks and fine-tuning them for binding site prediction [11]. This approach benefits from the knowledge acquired in the pretraining phase. Evaluating the performance of prediction methods is crucial. Researchers design benchmark datasets, establish evaluation metrics (such as sensitivity, specificity, and AUC), and compare different algorithms to assess their effectiveness.

The information needed to understand protein structures is all contained in the protein sequence. However, extracting the structure from the sequence only is a difficult and time-consuming task. Consequently, structure-only-based models consistently outperform sequence-based models in terms of performance due to the availing of complete structures. Structure-based models, however, need precise protein structures as input to assure model performance. On the other hand, because feature extraction frequently relies on manual design and does not produce a refined initial representation, the performance of existing sequence-based models is still insufficient for practical application. Therefore, it is imperative to create an end-to-end model without the use of handcrafted features. In this work, we identify several limitations to the work done in [20] and propose alternate solutions to overcome the limitations. More specifically, our contributions are the following:

1. It is a well-established fact in the literature that the neural network-based methods do not work efficiently as compared to simple Machine Learning (ML) classifiers (e.g. tree-based methods) in the case of tabular data [10,16,21]. Therefore, instead of using a One-dimensional Convolutional Neural Network (1DCNN) for the underlying classification (and including all discussion around that) as done in [20], we use simple ML classifiers for the underlying supervised analysis.
2. Authors in [20] use ProtBert [9] as the pre-trained model to generate the embeddings for each amino acid within protein sequences. We replace that with a more efficient SeqVec [13] pre-trained model. The choice of replacing ProtBert with SeqVec is due to its demonstrated effectiveness in learning relevant features for our task (i.e. Binding site prediction).
3. We propose a lightweight model using the idea of Sparse Coding, which combines the power of *k*-mers and one-hot encoding to design efficient initial embeddings for the amino acids. The only parameter in this sparse coding-based embedding method is *k* (contextual window size for amino acids), which is significantly lesser compared to complex models like ProtBert and SeqVec. This *Almost Parameter Free* approach makes Sparse coding an ideal choice for *fast* binding site prediction.

## 2 Related Work

The prediction of protein-DNA binding sites is a critical task in computational biology, with applications ranging from understanding gene regulation to designing novel therapeutic agents [6]. Over the years, various computational methods have been developed to tackle this complex problem, each leveraging different techniques and approaches. In this section, we review the key literature in the field of protein-DNA binding site prediction, focusing on different methodologies, challenges, and advancements. Evolutionary information has been a cornerstone of protein-DNA binding site prediction [33]. Methods utilizing multiple sequence alignments (MSAs) [1,41] and phylogenetic profiles [17] have shown promising results. Techniques like DRNAPred [42] and DNAPred [49] incorporate evolutionary conservation patterns to identify potential binding sites. SVMnuc [34] and NCBRPred [44] also utilize evolutionary information for distinguishing binding sites. However, most of these existing methods are computationally expensive, or not showing state of the art performance.

Traditional machine-learning techniques have been extensively used in the context of binding site prediction. Methods like SVMnuc [34] and DBPred [27] incorporate support vector machines (SVMs) to classify binding sites based on a set of engineered features derived from sequence and structure data. These methods have demonstrated reasonable predictive performance and often rely on well-curated training datasets. However, these methods may not be able to scale effectively to bigger datasets.

Recent advancements in deep learning have led to the development of more complex models for protein-DNA binding site prediction [33] and protein sequence analysis [25,24,22,23]. ProtBert [9]. A pre-trained transformer model adapted from natural language processing has shown its potential to capture intricate sequence patterns. The combination of ProtBert with 1D convolutional neural networks (1DCNN) has been explored to enhance performance in identifying binding sites. However, it is not clear if these methods can be generalized on other biological domains effectively.

Transfer learning from related domains, such as language models, has become a prominent technique [26]. Pre-trained models like ProtBert and SeqVec [13], inspired by NLP models, have shown success in capturing high-level features in protein sequences. These models provide a foundation for building more specialized predictors with fewer labeled samples. However, such models could be domain-specific and may not generalize for different tasks. The authors in [3] combined structure-based and sequence methods for protein analyses. SeqVec introduces embeddings that capture the biophysical properties of protein sequences by training on vast unlabeled protein data. These embeddings, derived from a language model, have demonstrated their potential in improving predictions. Sparse coding techniques, which do not require labeled data, have also been explored to generate embeddings that preserve important context [38].

## 3 Proposed Approach

We propose a protein-ligand binding sites prediction framework to perform the binding site prediction of a given protein sequence. The overall architecture of the proposed model comprised two main modules: the sequence embedding module and the classification module.

### 3.1 Sequence Embedding Module

The sequence embedding module leverages two distinct techniques, namely SeqVec and Sparse Coding, to create fixed-length embeddings for individual amino acids within a protein sequence.

*SeqVec [13]* It is a pre-trained protein language model that captures intricate sequence patterns and semantic information inherent to protein sequences. It is based on Embeddings from Language Models (ELMo) [32], commonly applied in natural language processing to create continuous vector representations (embeddings) for protein sequences. These embeddings, named SeqVec (Sequence-to-Vector), capture biophysical properties from unlabeled data (UniRef50) and enable simple neural networks to excel in various tasks. The key innovation of SeqVec lies in its use of ELMo to capture the biophysical properties of protein sequences. ELMo’s ability to learn contextualized embeddings from unlabeled protein sequences enables SeqVec to generate embeddings that encode relevant information about protein structure and function. This approach offers an alternative to the traditional use of evolutionary information and provides a scalable solution for analyzing protein sequences, particularly in scenarios involving large-scale proteomics data. Note that we justify the preference for SeqVec over other protein-based pre-trained language models, such as ProtBert [9] (as used in CLAPE [20]) due to its demonstrated effectiveness in learning relevant features for our task.

The architecture of SeqVec contains the following steps:

#### 1. ELMo Pre-training

ELMo, originally designed for natural language processing, is a bi-directional language model that learns to predict the likelihood of the next word in a sentence given the surrounding words. It does so by training on massive amounts of unlabeled text data, such as Wikipedia articles. ELMo develops contextualized embeddings that capture the syntax and semantics of the language. In the context of SeqVec, ELMo is trained on large protein sequence databases, specifically UniRef50, to predict the next amino acid in a sequence based on its neighboring amino acids.

#### 2. Embedding Extraction

After pre-training, ELMo produces embeddings for amino acids in a protein sequence. These embeddings capture the contextual information about each amino acid based on its surrounding amino acids in the sequence.

#### 3. Sequence Embeddings

The output of ELMo for each amino acid is a continuous vector representation that captures the biophysical properties of the protein sequence. These embeddings are referred to as SeqVec embeddings and serve as the representation of the protein sequence.

#### 4. Embeddings for Prediction Tasks

The SeqVec embeddings can be used as features for various protein prediction tasks, such as secondary structure prediction, intrinsic disorder prediction, subcellular localization prediction, and more. These embeddings are fed into neural networks or other machine learning models to perform these tasks.

*Sparse Coding* Since tunning a language model could still be expensive and it may not generalize better, we proposed a sparse coding-based alternative, which involves the power of *k*-mers (for neighborhood context capturing) and one-hot encoding (for generic embedding generation) to transform amino acids into numerical representations. The utilization of Sparse Coding is justifiable by its ability to capture local compositional information within amino acids of the protein sequences, enhancing our model’s capability to learn meaningful patterns associated with binding sites. For this purpose, we take a *k*-mer (where *k* = 9, which is decided using the standard validation set approach) as a sliding window for each amino acid. Then we design a one-hot encoding-based representation for the *k*-mer, which acts as the local embedding for the given amino acid. In this way, we design embedding for each amino acid, which is then used as input for supervised analysis using machine learning and deep learning models.

One exception occurs in our sparse coding-based embedding when the sliding window (*k*-mer) reaches the end of the sequence. In that case, for the remaining *n − k* amino acids (where *n* is the protein sequence length), we take the *k*-mers-based sliding window in reverse order and repeat the one-hot encoding step, hence preserving the neighborhood context. The overall workflow of the Sparse Coding embedding generation technique is illustrated in Figure 1.

**Fig. 1:**
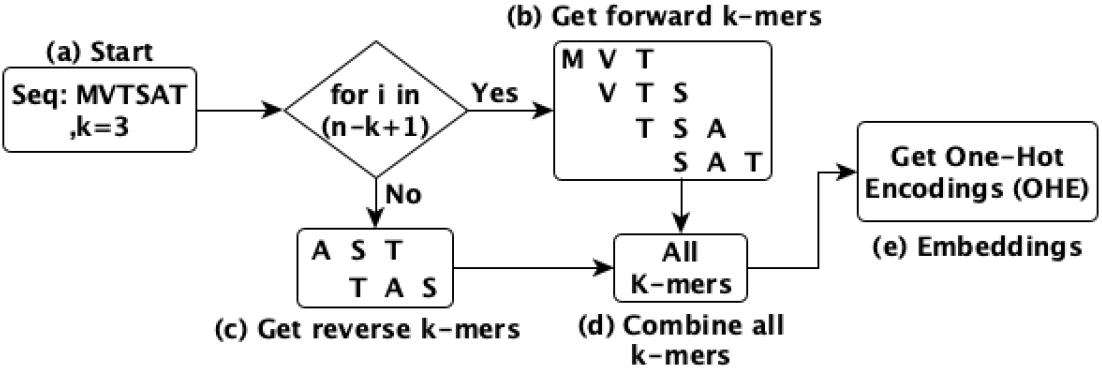
Workflow of Sparse Coding embedding generation method for a given sequence.

### 3.2 Classification Module

After designing the embeddings for each amino acid within the protein sequence to bind site prediction, the next step is to select efficient classification models to perform the actual site prediction. For this purpose, the features generated by the sequence embedding module (i.e. SeqVec and Sparse Coding) are fed into the classification module, which is composed of multiple machine-learning classifiers. For the same binding site prediction problem, authors in [20] propose the use of a four “one-dimensional convolutional neural network” (1DCNN) model as the backbone network. The raw dimension of the input is 1024, and the output dimensions of the four layers are 1024, 128, 64, and 2, respectively. Each layer has a stride of 1 and is followed by a batch normalization layer (except for the last one). The layers are also accompanied by varying sizes of kernel filters and paddings. The kernel sizes are 7, 5, 3, 5, and the paddings are 3, 2, 1, 2.

The 1DCNN is designed to capture neighboring information within protein sequences and employs operations like max pooling for down-sampling. Padding is applied for different convolutional kernel sizes to maintain the same sequence length for input and output features, ensuring a unified token-level classification outcome. The activation function ReLU (rectified linear unit) introduces nonlinearity to the model, and techniques such as dropout and batch normalization are utilized to enhance model robustness and generalization. The classification head is an integral part of the model, employing a Softmax function. This function scales the output values between 0 and 1, representing mutually exclusive prediction scores. These scores reflect the probability of a given residue being a DNA-binding site. The classification head is then used to predict DNA-binding sites within protein sequences.

While deep learning model, such as 1DCNN, exhibits remarkable capacity in various tasks, the dataset size, task complexity, and interpretability considerations have guided our choice towards machine-learning classifiers (i.e. Naive Bayes, Multi-Layer Perceptron, K-Nearest Neighbors, Random Forest, Logistic Regression, and Decision Tree). These classifiers collectively analyze the encoded features and make predictions about the presence of ligand binding sites in the protein sequence. Moreover, it is well known in the literature that the neural network-based methods do not perform optimally as compared to simple Machine Learning (ML) classifiers (e.g. tree-based methods) in the case of tabular data [10,16,21]. Therefore, we decided to use simple ML models for the downstream supervise analysis (i.e. binding site prediction), as done in the literature [4,2,36].

By integrating these modules, our proposed framework strives to provide accurate predictions of protein-ligand binding sites, leveraging the strengths of SeqVec and Sparse Coding for feature representation, and harnessing machine-learning classifiers for classification tasks. This design rationale ensures a well-rounded approach to predicting binding probabilities while considering the intricacies of the protein-ligand interaction problem.

To demonstrate the power of simple ML models over the deep learning models, we fine-tuned the existing ProtBert [9] model (as used in CLAPE [20]) to generate embeddings for the amino acids and performed binding site predictions as well. The finetuning hyper-parameters are ADAM optimizer, 25 batch size, and 10 training epochs. A loss function is formed by combining the focal loss [19] and triplet center loss (TCL) [12] to handle the data imbalance issue effectively, and it’s defined as *Loss* = *L*_*focal*_ + *λL*_*tcl*_, where *λ* is a hyperparameter with 0.1 value.

## 4 Experimental Setup

This section discusses the details of the datasets used for conducting the experiments along with the employed evaluation metrics and baseline methods. All experiments are conducted using a server having Intel(R) Xeon(R) CPU E7-4850 v4 @ 2.40*GHz* with Ubuntu 64 bit OS (16.04.7 LTS Xenial Xerus) having 3023 GB memory. Our code is available online ^1^.

### Dataset Statistics

We perform the binding site classification task using DNA-based datasets. These datasets were preprocessed to improve the model’s robustness and avoid data imbalance bias. In both datasets, the binding sites were defined as residues with a distance of *<* 0.5 (threshold value) +*R*, where *R* represents the sum of the Van der Waals radius of the two nearest atoms between the residue and the nucleic acid molecule. In each dataset, protein sequences are present along with their respective binding site indication as labels. For instance, a sample sequence from the train set of Dataset1 is “ARRIGHPYQNRTPPKRKK” where the alphabets represent the amino acids of the respective protein sequence and the labels are “001111110011011100”. These labels indicate if the corresponding amino acid has DNA binding site capacity or not i.e. 1 is for the binding site & 0 for the non-binding site. The details of these data are given in Table 1.

**Table 1:** The details of maximum, minimum & average lengths of sequences in the datasets 1 & 2 respectively. The number of binding and non-binding sites present in the test and train sets of each DNA-based dataset respectively [20]

|  |  | Length Statistics |  |  |  |  |  |
| --- | --- | --- | --- | --- | --- | --- | --- |
|  |  | Count | Max | Min | Avg | Binding | Non-Binding |
| Dataset 1 [27,31,45] | Train | 646 | 1937 | 36 | 279.02 | 15636 | 298503 |
|  | Test | 46 | 743 | 62 | 236.43 | 965 | 9911 |
| Dataset 2 [39,43,20] | Train | 573 | 3969 | 45 | 486.28 | 14479 | 145404 |
|  | Test | 129 | 968 | 55 | 290.81 | 2240 | 35275 |

### Evaluation Metrics

To evaluate the performance of the binding site prediction task, we used various evaluation metrics. The metrics are specificity, precision, recall, F1-score, ROC AUC, and Matthews correlation coefficient (MCC). We have reported the average score for each metric after 5 runs.

### Baseline Models

We have compared the performance of our proposed methods with various baselines described in Table 2.

**Table 2:** The baseline methods used to evaluate the system.

| Baseline | Description |
| --- | --- |
| DRNAPred [42] | A sequence-based method for predicting and differentiating between DNA- and RNA-binding residues. |
| DNAPred [49] | It introduces a novel two-stage imbalanced learning algorithm called Ensembled Hyperplane-Distance-Based Support Vector Machines for the prediction of protein-DNA binding sites. |
| SVMnuc [34] | It is an ab-initio method devised to predict nucleic acids-binding residues, addressing the challenge of accurately identifying these biologically significant sites. |
| NCBRPred [44] | It is designed to predict nucleic acid binding residues within proteins. NCBRPred adopts a multilabel sequence labeling model (MSLM). |
| DBPred [27] | A server-based tool for predicting DNA-binding residues in proteins. The sequence module allows to predict the DNA-interacting residues using the binary and physicochemical profiles. |
| CLAPE [20] | This approach effectively combines the power of a pre-trained protein language model (i.e. ProBert) with contrastive learning techniques (i.e. 1DCNN model) to do prediction of DNA binding residues in proteins |

### t-SNE Visualization

A popular visualization technique, named t-SNE [37], is employed by us for visualization.

The t-SNEs against Dataset1 for the top 3 longest sequences are depicted in Figure 2. We can observe that in all the plots the clusters are overlapping and non-definite.

**Fig. 2:**
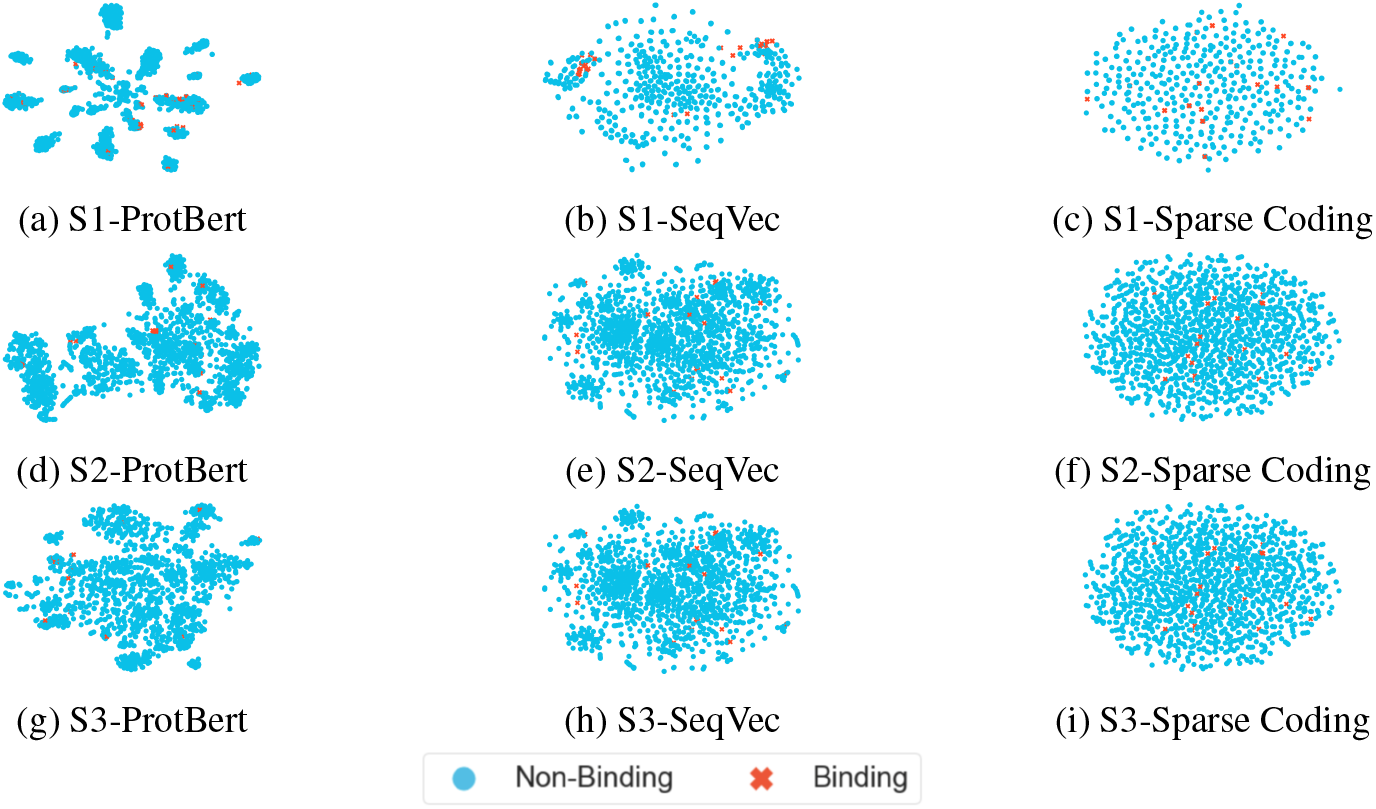
t-SNE visualization of embeddings generated by different embedding generation methods (ProtBert, SeqVec, & Sparse Coding) using top 3 longest sequences (S1, S2, S3) from **Dataset 1**. The Figure is best seen in color.

The binding instances are less visible and scattered throughout the plots in each figure, and a reason for this could be the data imbalance issue in the dataset i.e. number of binding instances is much less than the non-binding ones. Moreover, the Sparse Coding technique is yielding very similar cluster structures for all three sequences, while Prot-Bert and SeqVec show some variation in the structures. Overall, the patterns illustrate that none of the embedding methods for any sequence can generate very clear clusters for both the binding and non-binding classes in a 2-dimensional space.

The t-SNE visualization of the top 3 longest sequences from Dataset2 are shown in Figure 3. We can observe that for any sequences against the ProtBert technique, the binding class instances are almost invisible. This indicates that this method can not preserve a good structure for less frequent classes from the dataset in a 2-dimensional space. Furthermore, the SeqVec and Sparse Coding mechanisms illustrate binding clusters being scattered across the plots for all the sequences. Overall, yet again no definite and clear cluster structures can be viewed for Dataset2 and it can also be due to the class imbalance challenge.

**Fig. 3:**
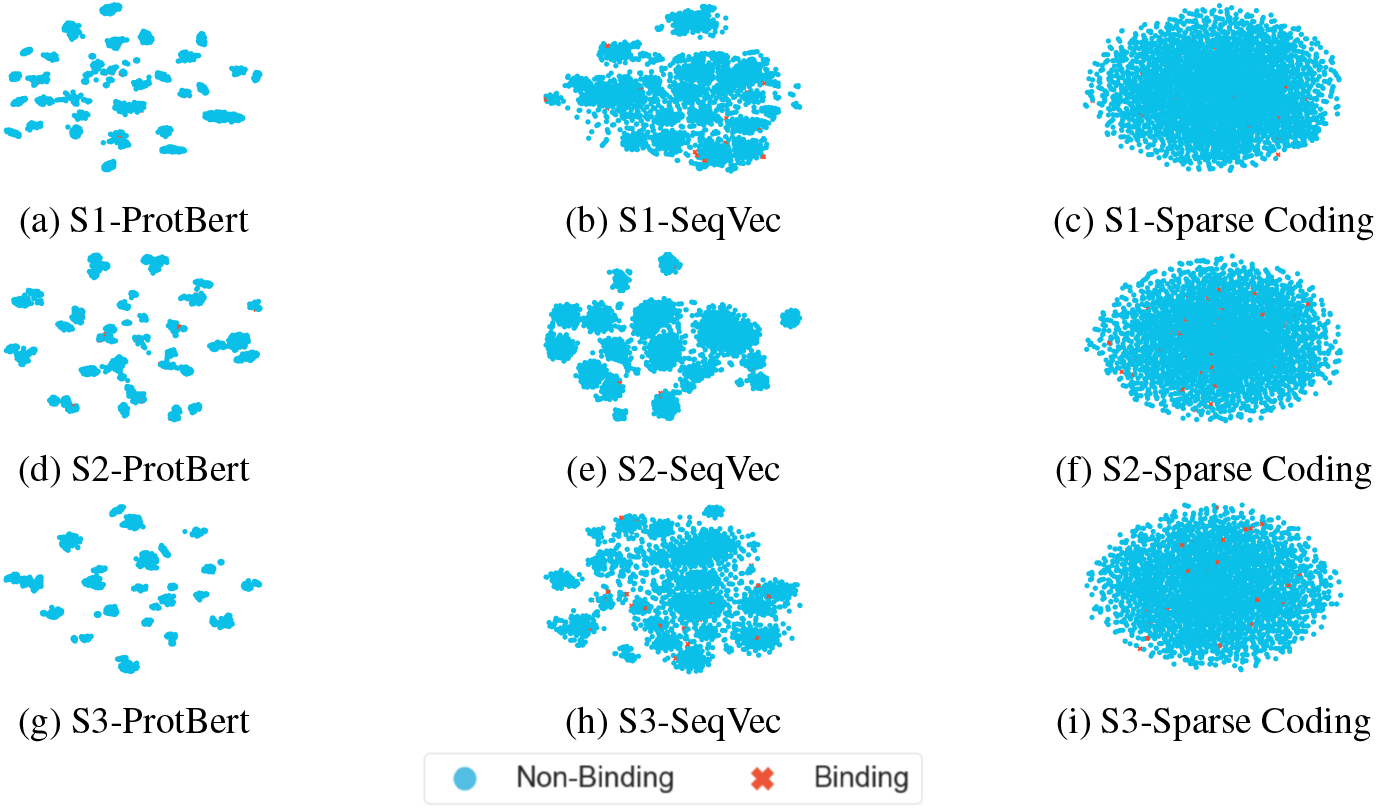
t-SNE visualization of embeddings generated by different embedding generation methods (ProtBert, SeqVec, & Sparse Coding) using top 3 longest sequences (S1, S2, S3) from **Dataset 2**. The Figure is best seen in color.

## 5 Results And Discussion

The classification results for the proposed method and its comparison with the baselines are shown in Table 3 for Dataset 1. Compared to the baselines such as DRNAPred, DNAPred, SVMnuc, NCBRPred, DBPred, and ProtBert + 1DCNN, we can observe that ProtBert + ML classifiers (our pre-trained model) i.e. Naive Bayes, Multi-layer Perceptron, K-Nearest Neighbors, Random Forest, Logistic Regression, and Decision Tree, show near-perfect specificity and precision scores. This eventually means that the number of correctly predicted non-binding sites is higher. Moreover, the accuracy of the residues predicted as DNA-binding sites is also higher. However, for Recall and ROC-AUC, the baseline NCBRPred shows higher performance while ProtBert + 1DCNN shows superior performance in the case of F1 and MCC scores. More complex models like ProtBert + 1DCNN may have a higher capacity to capture intricate patterns in the data, which could lead to better F1 and MCC scores. For our SeqVec + ML classifiers and Sparse coding + ML/DL classifiers, we can again observe a near-perfect specificity score. One interesting insight to note here is that since the Sparse coding-based embedding method is completely unsupervised and does not involve any expensive model training, it is still able to achieve a higher specificity score. This is due to the fact that it preserves the neighborhood context efficiently within the generated embeddings. The reason for the simpler models to excel in specific metrics, such as ProtBert + ML classifiers achieving high specificity and precision due to their focused decision boundaries.

**Table 3:** Binding site prediction (classification) results for different evaluation metrics using the proposed and baseline methods for **Dataset 1**. The best values are shown in bold. Dashes “-” in the model column mean they were end-to-end models and used as described in respective original studies

| Method | Model | Spec. | Prec. | Recall | F1 (Bina.) | ROC-AUC | MCC |
| --- | --- | --- | --- | --- | --- | --- | --- |
| DRNAPred | - | 0.692 | 0.185 | 0.677 | 0.291 | 0.755 | 0.226 |
| DNAPred | - | 0.655 | 0.157 | 0.671 | 0.254 | 0.730 | 0.194 |
| SVMnuc | - | 0.666 | 0.154 | 0.668 | 0.250 | 0.715 | 0.192 |
| NCBRPred | - | 0.674 | 0.165 | 0.677 | 0.265 | 0.713 | 0.207 |
| DBPred | - | 0.784 | 0.243 | **0.708** | 0.362 | **0.794** | 0.320 |
| ProBert | 1DCNN | 0.834 | 0.307 | 0.658 | **0.380** | 0.746 | **0.339** |
|  | NB | 0.775 | 0.167 | 0.464 | 0.246 | 0.619 | 0.093 |
|  | MLP | 0.971 | 0.466 | 0.254 | 0.329 | 0.613 | 0.294 |
|  | KNN | 0.981 | 0.500 | 0.191 | 0.277 | 0.586 | 0.271 |
|  | RF | **0.999** | **0.999** | 0.002 | 0.004 | 0.501 | 0.043 |
|  | LR | 0.994 | 0.672 | 0.117 | 0.199 | 0.555 | 0.265 |
|  | DT | 0.942 | 0.211 | 0.159 | 0.182 | 0.550 | 0.096 |
| SeqVec | 1DCNN | 0.972 | 0.418 | 0.280 | 0.293 | 0.626 | 0.276 |
|  | NB | 0.807 | 0.178 | 0.431 | 0.252 | 0.619 | 0.103 |
|  | MLP | 0.963 | 0.379 | 0.231 | 0.287 | 0.597 | 0.228 |
|  | KNN | 0.991 | 0.692 | 0.191 | 0.300 | 0.591 | 0.356 |
|  | RF | **0.999** | 0.851 | 0.023 | 0.046 | 0.511 | 0.135 |
|  | LR | 0.997 | 0.715 | 0.075 | 0.136 | 0.536 | 0.219 |
|  | DT | 0.933 | 0.197 | 0.167 | 0.181 | 0.551 | 0.088 |
| Sparse Coding<br>(kmers+OHE) | 1DCNN | **0.999** | 0.000 | 0.000 | 0.000 | 0.500 | 0.000 |
|  | NB | 0.938 | 0.096 | 0.067 | 0.079 | 0.503 | 0.007 |
|  | MLP | **0.999** | 0.000 | 0.000 | 0.000 | 0.500 | 0.000 |
|  | KNN | 0.997 | 0.115 | 0.003 | 0.006 | 0.500 | 0.009 |
|  | RF | 0.997 | 0.289 | 0.011 | 0.021 | 0.504 | 0.041 |
|  | LR | **0.999** | 0.000 | 0.000 | 0.000 | 0.500 | 0.000 |
|  | DT | 0.944 | 0.102 | 0.065 | 0.079 | 0.507 | 0.011 |

The classification results for the proposed method and its comparison with the baselines are shown in Table 4 for Dataset 2. We can observe that for the specificity, precision, and MCC, both pre-trained models (i.e. ProBert and SeqVec) and the Sparse coding-based method show higher scores using simple ML classifiers rather than using comparatively more complex 1DCNN model. For F1 and ROC-AUC, we can observe that DNAPred performs the best.

**Table 4:** Binding site prediction (classification) results for different evaluation metrics using the proposed and baseline methods for **Dataset 2**. The best values are shown in bold. Dashes “-” in the model column mean they were end-to-end models and used as described in respective original studies.

| Method | Model | Spec. | Prec. | Recall | F1 (Bina.) | ROC-AUC | MCC |
| --- | --- | --- | --- | --- | --- | --- | --- |
| DRNAPred | - | 0.937 | 0.190 | 0.233 | 0.210 | 0.693 | 0.155 |
| DNAPred | - | 0.954 | 0.353 | 0.396 | **0.373** | **0.845** | 0.332 |
| SVMnuc | - | 0.966 | 0.371 | 0.316 | 0.341 | 0.812 | 0.304 |
| NCBRPred | - | 0.969 | 0.312 | 0.392 | 0.347 | 0.823 | 0.313 |
| ProBert | 1DCNN | 0.830 | 0.242 | **0.619** | 0.317 | 0.725 | 0.221 |
|  | NB | 0.761 | 0.141 | 0.618 | 0.230 | 0.690 | 0.100 |
|  | MLP | 0.954 | 0.305 | 0.318 | 0.311 | 0.636 | 0.225 |
|  | KNN | 0.974 | 0.310 | 0.179 | 0.227 | 0.577 | 0.181 |
|  | RF | **0.999** | 0.545 | 0.002 | 0.005 | 0.501 | 0.035 |
|  | LR | 0.979 | 0.489 | 0.305 | 0.376 | 0.642 | **0.354** |
|  | DT | 0.910 | 0.123 | 0.198 | 0.157 | 0.554 | 0.059 |
| SeqVec | 1DCNN | 0.960 | 0.328 | 0.287 | 0.283 | 0.623 | 0.292 |
|  | NB | 0.753 | 0.089 | 0.382 | 0.145 | 0.568 | 0.035 |
|  | MLP | 0.954 | 0.262 | 0.253 | 0.257 | 0.604 | 0.175 |
|  | KNN | 0.986 | 0.503 | 0.215 | 0.301 | 0.601 | 0.303 |
|  | RF | **0.999** | **0.782** | 0.051 | 0.096 | 0.525 | 0.194 |
|  | LR | 0.991 | 0.512 | 0.146 | 0.227 | 0.568 | 0.252 |
|  | DT | 0.882 | 0.107 | 0.222 | 0.144 | 0.552 | 0.046 |
| Sparse Coding<br>(kmers+OHE) | 1DCNN | **0.999** | 0.000 | 0.000 | 0.000 | 0.500 | 0.000 |
|  | NB | 0.986 | 0.058 | 0.012 | 0.021 | 0.499 | 0.000 |
|  | MLP | **0.999** | 0.103 | 0.001 | 0.002 | 0.500 | 0.005 |
|  | KNN | 0.993 | 0.066 | 0.007 | 0.012 | 0.500 | 0.002 |
|  | RF | 0.997 | 0.109 | 0.004 | 0.007 | 0.500 | 0.009 |
|  | LR | **0.999** | 0.000 | 0.000 | 0.000 | 0.500 | 0.000 |
|  | DT | 0.900 | 0.062 | 0.104 | 0.078 | 0.502 | 0.002 |

We utilized a deep language model, ProteinBert [5], to get the feature embeddings of the protein sequences from our datasets and employed those embeddings to perform DNA binding site classification using our classification models (given in Section 3.2). ProteinBert is designed explicitly for protein sequences and it consists of both local and global representations. We obtain the global representation in our experiments.

The DNA-binding site classification results of Dataset 1 are given in Table 5. The results illustrate that the NB classifier depicts maximum performance in terms of recall, f1 score, and ROC-AUC metrics, while MCC & precision is optimal for the RF classifier and specificity for MLP. However, overall ProteinBert is not showing optimal performance as compared to the other methods (mentioned in Table 3).

**Table 5:** Binding site prediction (classification) results for different evaluation metrics using the ProteinBert embedding generation method for **Dataset 1**. The best values are shown in bold.

| Method | Model | Spec. | Prec. | Recall | F1 (Bina.) | ROC-AUC | MCC |
| --- | --- | --- | --- | --- | --- | --- | --- |
| Protein<br>Bert | 1DCNN | 0.998 | 0.000 | 0.044 | 0.000 | 0.521 | 0.060 |
|  | NB | 0.529 | 0.106 | **0.574** | **0.179** | **0.551** | 0.025 |
|  | MLP | **0.999** | 0.000 | 0.000 | 0.000 | 0.500 | 0.000 |
|  | KNN | 0.936 | 0.154 | 0.119 | 0.134 | 0.527 | 0.050 |
|  | RF | 0.972 | **0.215** | 0.078 | 0.115 | 0.525 | **0.074** |
|  | LR | 0.998 | 0.000 | 0.000 | 0.000 | 0.500 | 0.000 |
|  | DT | 0.967 | 0.181 | 0.073 | 0.104 | 0.520 | 0.055 |

Moreover, the classification results obtained from Dataset 2 are reported in Table 6. We can observe that the NB model is outperforming others in terms of recall, f1 score, and ROC-AUC metrics. Precision and MCC have the highest values against the DT model, while specificity is optimal for the MLP classifier. However, yet again the ProteinBert is unable to achieve optimal performance as compared to the other methods (mentioned in Table 4).

**Table 6:** Binding site prediction (classification) results for different evaluation metrics using the ProteinBert embedding generation method for **Dataset 2**. The best values are shown in bold.

| Method | Model | Spec. | Prec. | Recall | F1 (Bina.) | ROC-AUC | MCC |
| --- | --- | --- | --- | --- | --- | --- | --- |
| Protein<br>Bert | 1DCNN | 0.998 | 0.000 | 0.002 | 0.000 | 0.510 | 0.000 |
|  | NB | 0.724 | 0.119 | **0.536** | **0.194** | **0.630** | 0.067 |
|  | MLP | **0.999** | 0.000 | 0.004 | 0.000 | 0.500 | 0.000 |
|  | KNN | 0.979 | 0.068 | 0.021 | 0.032 | 0.500 | 0.001 |
|  | RF | 0.996 | 0.230 | 0.016 | 0.029 | 0.506 | 0.044 |
|  | LR | 0.998 | 0.002 | 0.003 | 0.000 | 0.500 | 0.010 |
|  | DT | 0.991 | **0.241** | 0.040 | 0.068 | 0.515 | **0.071** |

## 6 Conclusion

In this study, we have addressed the challenging problem of binding site prediction using an innovative and comprehensive approach. Our work capitalizes on the syn-ergy between conventional bioinformatics techniques, state-of-the-art protein language models, and advanced machine learning classifiers. Through meticulous experimentation and rigorous evaluation, we have demonstrated the superiority of our proposed approach over existing models. Our model exhibits robust predictive behavior as evidenced by higher predictive values on benchmark datasets. The flexibility and generalization capacity of our models is highlighted by their adaptability as a universal framework for binding site prediction across diverse protein-ligand binding scenarios. By leveraging SeqVec, a powerful pre-trained model, we capture intricate sequence features effectively. Additionally, we propose a lightweight model based on Sparse Coding, which combines *k*-mers and one-hot encoding to generate efficient initial embeddings. This approach’s parameter efficiency positions it as a promising candidate for rapid binding site prediction. As we continue to refine and expand our approach, we envision its potential to drive breakthroughs across various domains of biology and genetics. Exploring the integration of epigenetic information and investigating ensemble methods to combine predictions from diverse models could enhance the performance of binding site prediction, thus advancing our understanding of intricate cellular processes.

## 7 Acknowldgement

Sarwan Ali, Prakash Chourasia, and Murray Patterson would like to thank Molecular Basis of Disease (MBD) and Georgia State University for their support and resources.

## Footnotes

1 https://github.com/sarwanpasha/Binding-Site-Prediction

## References

1. Ahmad, S., and Sarai, A. Pssm-Based Prediction Of Dna Binding Sites In Proteins. Bmc Bioinformatics 6 (2005), 1–6.

2. Ali, S., Ali, T. E., Murad, T., Mansoor, H., and Patterson, M. Molecular Se-Quence Classification Using Efficient Kernel Based Embedding. Information Sciences 679 (2024), 121100.

3. Ali, S., Chourasia, P., and Patterson, M. Pdb2vec: Using 3d Structural Information For Improved Protein Analysis. In Isbra (2023), Pp. 376–386.

4. Ali, S., Shabbir, M., Mansoor, H., Chourasia, P., and Patterson, M. Elliptic Geometry-Based Kernel Matrix For Improved Biological Sequence Classification. Knowledge-Based Systems (2024), 112479.

5. Brandes, N., Ofer, D., et al. Proteinbert: A Universal Deep-Learning Model Of Protein Sequence and Function. Bioinformatics 38, 8 (2022), 2102–2110.

6. Collie, G. W., and Parkinson, G. N. The Application Of Dna and Rna G-Quadruplexes To Therapeutic Medicines. Chemical Society Reviews 40, 12 (2011), 5867–5892.

7. Dey, B., Thukral, S., Krishnan, S. A., Chakrobarty, M., Gupta, S., Manghani, C., and Rani, V. Dna–Protein Interactions: Methods For Detection and Analysis. Molecular And Cellular Biochemistry 365 (2012), 279–299.

8. Echols, H. Multiple Dna-Protein Interactions Governing High-Precision Dna Transactions. Science 233 4768 (1986), 1050–6.

9. Elnaggar, A., Heinzinger, M., et al. Prottrans: Towards Cracking The Language Of Life’S Code Through Self-Supervised Learning. Biorxiv (2021).

10. Grinsztajn, L., Oyallon, E., and Varoquaux, G. Why Do Tree-Based Models Still Outperform Deep Learning On Tabular Data? Arxiv Preprint Arxiv:2207.08815 (2022).

11. Han, W., Pang, B., and Wu, Y. N. Robust Transfer Learning With Pretrained Language Models Through Adapters. Arxiv Abs/2108.02340 (2021).

12. He, X., Zhou, Y., Zhou, Z., Bai, S., and Bai, X. Triplet-Center Loss For Multi-View 3d Object Retrieval. In Proceedings Of The Ieee Conference On Computer Vision And Pattern Recognition (2018), Pp. 1945–1954.

13. Heinzinger, M., Elnaggar, A., et al. Modeling Aspects Of The Language Of Life Through Transfer-Learning Protein Sequences. Bmc Bioinformatics 20, 1 (2019), 1–17.

14. Ivanciuc, O., Oezguen, N., Mathura, V. S., Schein, C. H., Xu, Y., and Braun, W. Using Property Based Sequence Motifs and 3d Modeling To Determine Structure and Functional Regions Of Proteins. Current Medicinal Chemistry 11 5 (2004), 583–93.

15. Jones, S., Van Heyningen, P., Berman, H. M., and Thornton, J. M. Protein-Rna Interactions: A Structural Analysis. Nucleic Acids Research 29 4 (2001), 943–54.

16. Joseph, M., and Raj, H. Gate: Gated Additive Tree Ensemble For Tabular Classification and Regression. Arxiv Preprint Arxiv:2207.08548 (2022).

17. La, D., and Kihara, D. A Novel Method For Protein–Protein Interaction Site Prediction Using Phylogenetic Substitution Models. Proteins: Structure, Function, And Bioinformatics 80, 1 (2012), 126–141.

18. Lichtarge, O., Bourne, H. R., et al. An Evolutionary Trace Method Defines Binding Surfaces Common To Protein Families. Journal Of Molecular Biology 257 2 (1996), 342–58.

19. Lin, T.-Y., Goyal, P., Girshick, R., He, K., and Dollár, P. Focal Loss For Dense Object Detection. In Proceedings Of The Ieee International Conference On Computer Vision (2017), Pp. 2980–2988.

20. Liu, Y., and Tian, B. Protein-Dna Binding Sites Prediction Based On Pre-Trained Protein Language Model and Contrastive Learning. Arxiv Preprint Arxiv:2306.15912 (2023).

21. Malinin, A., Prokhorenkova, L., and Ustimenko, A. Uncertainty In Gradient Boost-Ing Via Ensembles. In International Conference On Learning Representations (Iclr) (2021).

22. Murad, T., Ali, S., Khan, I., and Patterson, M. Spike2cgr: An Efficient Method For Spike Sequence Classification Using Chaos Game Representation. Machine Learning 112, 10 (2023), 3633–3658.

23. Murad, T., Ali, S., and Patterson, M. A New Direction In Membranolytic Anticancer Peptides Classification: Combining Spaced K-Mers With Chaos Game Representation. Procedia Computer Science 222 (2023), 666–675.

24. Murad, T., Ali, S., and Patterson, M. Weighted Chaos Game Representation For Molecular Sequence Classification. In Pacific-Asia Conference On Knowledge Discovery And Data Mining (2024), Pp. 234–245.

25. Murad, T., Chourasia, P., Ali, S., and Patterson, M. Dance: Deep Learning-Assisted Analysis Of Protein Sequences Using Chaos Enhanced Kaleidoscopic Images. Arxiv Preprint Arxiv:2409.06694 (2024).

26. Novakovsky, G., Saraswat, M., et al. Biologically Relevant Transfer Learning Improves Transcription Factor Binding Prediction. Genome Biology 22, 1 (2021), 1–25.

27. Patiyal, S., et al. A Deep Learning-Based Method For The Prediction Of Dna Interacting Residues In A Protein. Briefings In Bioinformatics 23, 5 (2022), Bbac322.

28. Pique-Regi, R., Degner, J. F., Pai, A. A., Gaffney, D. J., Gilad, Y., and Pritchard, J. K. Accurate Inference Of Transcription Factor Binding From Dna Sequence and Chromatin Accessibility Data. Genome Research 21 3 (2011), 447–55.

29. Polo, S. E., and Jackson, S. P. Dynamics Of Dna Damage Response Proteins At Dna Breaks: A Focus On Protein Modifications. Genes & Development 25 5 (2011), 409–33.

30. Ptashne, M. Gene Regulation By Proteins Acting Nearby and At A Distance. Nature 322 (1986), 697–701.

31. Qiu, J., Bernhofer, M., Heinzinger, M., et al. Prona2020 Predicts Protein–Dna, Protein–Rna, and Protein–Protein Binding Proteins and Residues From Sequence. Journal Of Molecular Biology 432, 7 (2020), 2428–2443.

32. Sarzynska-Wawer, J., Wawer, A., Pawlak, A., et al. Detecting Formal Thought Disorder By Deep Contextualized Word Representations. Psychiatry Research 304 (2021), 114135.

33. Si, J., Zhao, R., and Wu, R. An Overview Of The Prediction Of Protein Dna-Binding Sites. International Journal Of Molecular Sciences 16, 3 (2015), 5194–5215.

34. Su, H., Liu, M., Sun, S., Peng, Z., and Yang, J. Improving The Prediction Of Protein– Nucleic Acids Binding Residues Via Multiple Sequence Profiles and The Consensus Of Complementary Methods. Bioinformatics 35, 6 (2019), 930–936.

35. Tatarinova, T. V., Chekalin, E., Nikolsky, Y., Bruskin, S. A., Chebotarov, D., Mcnally, K. L., and Alexandrov, N. N. Nucleotide Diversity Analysis Highlights Functionally Important Genomic Regions. Scientific Reports 6 (2016).

36. Tayebi, Z., Ali, S., Murad, T., Khan, I., and Patterson, M. Pseaac2vec Protein Encoding For Tcr Protein Sequence Classification. Computers In Biology And Medicine 170 (2024), 107956.

37. Van Der M. L., and Hinton, G. Visualizing Data Using T-Sne. Journal Of Machine Learning Research (Jmlr) 9, 11 (2008).

38. Wu, J., Dong, Q., Gui, J., et al. Predicting Brain Amyloid Using Multivariate Morphometry Statistics, Sparse Coding, and Correntropy: Validation In 1,101 Individuals From The Adni and Oasis Databases. Frontiers In Neuroscience 15 (2021), 669595.

39. Xia, Y., Xia, C.-Q., Pan, X., and Shen, H.-B. Graphbind: Protein Structural Context Embedded Rules Learned By Hierarchical Graph Neural Networks For Recognizing Nucleic-Acid-Binding Residues. Nucleic Acids Research 49, 9 (2021), E51–E51.

40. Xu, D., Jalal, S. I., Sledge, G. W., and Meroueh, S. O. Small-Molecule Binding Sites To Explore Protein-Protein Interactions In The Cancer Proteome. Molecular Biosystems 12 10 (2016), 3067–87.

41. Yan, C., Terribilini, M., et al. Predicting Dna-Binding Sites Of Proteins From Amino Acid Sequence. Bmc Bioinformatics 7 (2006), 1–10.

42. Yan, J., et al. Drnapred, Fast Sequence-Based Method That Accurately Predicts and Discriminates Dna-And Rna-Binding Residues. Nucleic Acids Research 45, 10 (2017), E84–E84.

43. Yang, J., Roy, A., and Zhang, Y. Biolip: A Semi-Manually Curated Database For Biologically Relevant Ligand–Protein Interactions. Nucleic Acids Research 41 (2012), D1096–D1103.

44. Zhang, J., Chen, Q., and Liu, B. Ncbrpred: Predicting Nucleic Acid Binding Residues In Proteins Based On Multilabel Learning. Briefings In Bioinformatics 22, 5 (2021), Bbaa397.

45. Zhang, J., Ma, Z., and Kurgan, L. Comprehensive Review and Empirical Analysis Of Hallmarks Of Dna-, Rna-And Protein-Binding Residues In Protein Chains. Briefings In Bioinformatics 20, 4 (2019), 1250–1268.

46. Zhang, Q., Zhu, L., and Shuang Huang, D. High-Order Convolutional Neural Network Architecture For Predicting Dna-Protein Binding Sites. Ieee/Acm Transactions On Computational Biology And Bioinformatics 16 (2019), 1184–1192.

47. Zhao, J., Cao, Y., and Zhang, L. Exploring The Computational Methods For Proteinligand Binding Site Prediction. Computational And Structural Biotechnology Journal 18 (2020), 417–426.

48. Zhu, H., Wang, G., and Qian, J. Transcription Factors As Readers and Effectors Of Dna Methylation. Nature Reviews Genetics 17, 9 (2016), 551–565.

49. Zhu, Y.-H., Hu, J., Song, X.-N., and Yu, D.-J. Dnapred: Accurate Identification Of Dna-Binding Sites From Protein Sequence By Ensembled Hyperplane-Distance-Based Support Vector Machines. Journal Of Chemical Information And Modeling 59, 6 (2019), 3057–3071.

50. Śledź, P., and Caflisch, A. Protein Structure-Based Drug Design: From Docking To Molecular Dynamics. Current Opinion In Structural Biology 48 (2018), 93–102.

